# SiCTeC: an inexpensive, easily assembled Peltier device for rapid temperature shifting during single-cell imaging

**DOI:** 10.1101/2020.05.29.123166

**Authors:** Benjamin D. Knapp, Lillian Zhu, Kerwyn Casey Huang

**Author notes:** equal contributions (Benjamin D. Knapp, Lillian Zhu).

## Abstract

Single-cell imaging, combined with recent advances in image analysis and microfluidic technologies, have enabled fundamental discoveries of cellular responses to chemical perturbations that are often obscured by traditional liquid-culture experiments. Temperature is an environmental variable well known to impact growth and to elicit specific stress responses at extreme values; it is often used as a genetic tool to interrogate essential genes. However, the dynamic effects of temperature shifts have remained mostly unstudied at the single-cell level, due largely to engineering challenges related to sample stability, heatsink considerations, and temperature measurement and feedback. Additionally, the few commercially available temperature-control platforms are costly. Here, we report an inexpensive (<$110) and modular **Si**ngle-**C**ell **Te**mperature **C**ontroller (SiCTeC) device for microbial imaging, based on straightforward modifications of the typical slide-sample-coverslip approach to microbial imaging, that controls temperature using a ring-shaped Peltier module and microcontroller feedback. Through stable and precise (±0.15 °C) temperature control, SiCTeC achieves reproducible and fast (1-2 min) temperature transitions with programmable waveforms between room temperature and 45 °C with an air objective. At the device’s maximum temperature of 89 °C, SiCTeC revealed that *Escherichia coli* cells progressively shrink and lose cellular contents. During oscillations between 30 °C and 37 °C, cells rapidly adapted their response to temperature upshifts. Furthermore, SiCTeC enabled the discovery of rapid morphological changes and enhanced sensitivity to substrate stiffness during upshifts to nonpermissive temperatures in temperature-sensitive mutants of cell-wall synthesis enzymes. Overall, the simplicity and affordability of SiCTeC empowers future studies of the temperature dependence of single-cell physiology.

## Introduction

While chemical perturbations during single-cell imaging experiments have been made possible by microfluidic technologies [1, 2], other environmental variables such as temperature have been more difficult to precisely and rapidly manipulate during an experiment. Temperature has dramatic effects on virtually all cellular processes, including polymer behavior [3, 4], RNA and DNA polymerases [5, 6], ribosomal elongation [7], and overall enzyme kinetics and function [8]. These diverse, temperature-dependent processes have global impacts on cell growth, which cells must integrate and collectively optimize at each temperature [9].

Two predominant elements of experimental design limit our understanding of how cells respond to changes in temperature. First, bulk experiments are the standard for investigating the effects of temperature on steady-state cellular growth [10, 11]. By contrast, single-cell investigations reveal insights into morphological dynamics and population heterogeneity that cannot be achieved through bulk experimentation. Second, experimental designs tend to employ a single experimental temperature that maximizes growth rate (e.g. 37 °C for the enteric bacterium *Escherichia coli*). As a result, how cells respond to temperature fluctuations, such as during transitions into and out of a host, remains understudied; in *E. coli*, the few studies of temperature shifts have shown long time scales for growth-rate equilibration [12] and transient changes to the synthesis rate of tRNA synthetases [13]. While intriguing, these limited results highlight the need for a device that enables single-cell imaging to investigate how individual cells respond to temperature fluctuations at high spatiotemporal resolution.

Current microscope incubators are useful for maintaining temperature during time-lapse imaging, but are unable to precisely control the temperature of samples at short timescales. Consequently, a stage-top temperature controller for single-cell imaging is ideal for studying how cells transition between temperatures at high temporal resolution. Engineering such a device has remained challenging, as modifying temperature in real time requires sample stability, careful temperature measurement, and control feedback. Most commercial devices lack the capacity to program arbitrary temperature shift profiles and often can only heat samples to temperatures slightly higher than 37 °C. Further, the high cost of these devices (>$10,000) prohibits broad accessibility.

Recent low-cost alternatives to expensive laboratory techniques include electroporation [14], fluorescence microscopy and optogenetics [15], and nucleic acid extraction [16]. In addition to making science more open and accessible, particularly in resource-challenged institutions and countries [17], open-source experimentation can drive new applications that are not currently achievable with commercial equipment, akin to the impact of the development of the open-source Linux kernel on computing [18]. Inspired by these successes, here we built an affordable and flexible temperature controller for single-cell imaging that meets several crucial design objectives: (1) inexpensive and accessible components, (2) easy to assemble and compatible with traditional slide-mount techniques, (3) temperature-stable, (4) reusable, (5) capable of rapid temperature switching, (6) programmable complex temperature routines, and (7) digital temperature readout in real time. We provide instruction to construct this device in just a few hours using a ring-shaped Peltier module heater and open-source temperature control software for <$110. We used this device, the **Si**ngle-**C**ell **Te**mperature **C**ontroller (SiCTeC), to discover that wild-type *E. coli* cells rapidly adapt to temperature oscillations and lose cellular material at extreme temperatures, and that the phenotypes of temperature-sensitive *E. coli* cell-wall mutants quickly manifest but depend on the stiffness of the substrate on which they are grown. We anticipate that SiCTeC will drive fundamental research into the temperature dependence of cellular physiology and serve as a highly accessible device for precisely controlling environmental temperature.

## Results

### Design of the temperature-control device

To increase the utility of an inexpensive microscope stage-top temperature-control device, we based our development on existing imaging and sample preparation approaches. Our design modifies the traditional use of agarose hydrogel pads on glass slides as sample mounts [19] by adding a simple heating component and a temperature sensor (Fig. 1). To enable rapid and precise temperature changes, we implemented a Peltier-module design. Peltier modules are inexpensive semiconductor devices that drive large temperature gradients through the thermoelectric effect [20]. Although Peltier modules usually require large heatsinks to dissipate excess heat during cooling, we allowed the SiCTeC to self-heat (without a heatsink) to increase speed and performance, owing to its small size. SiCTeC consists of a ring-shaped Peltier module that permits illumination of samples during microscopy, which is crucial for single-cell investigations (Fig. 1A,B), and also ensures that heating is uniform across the sample.

**Figure 1:**
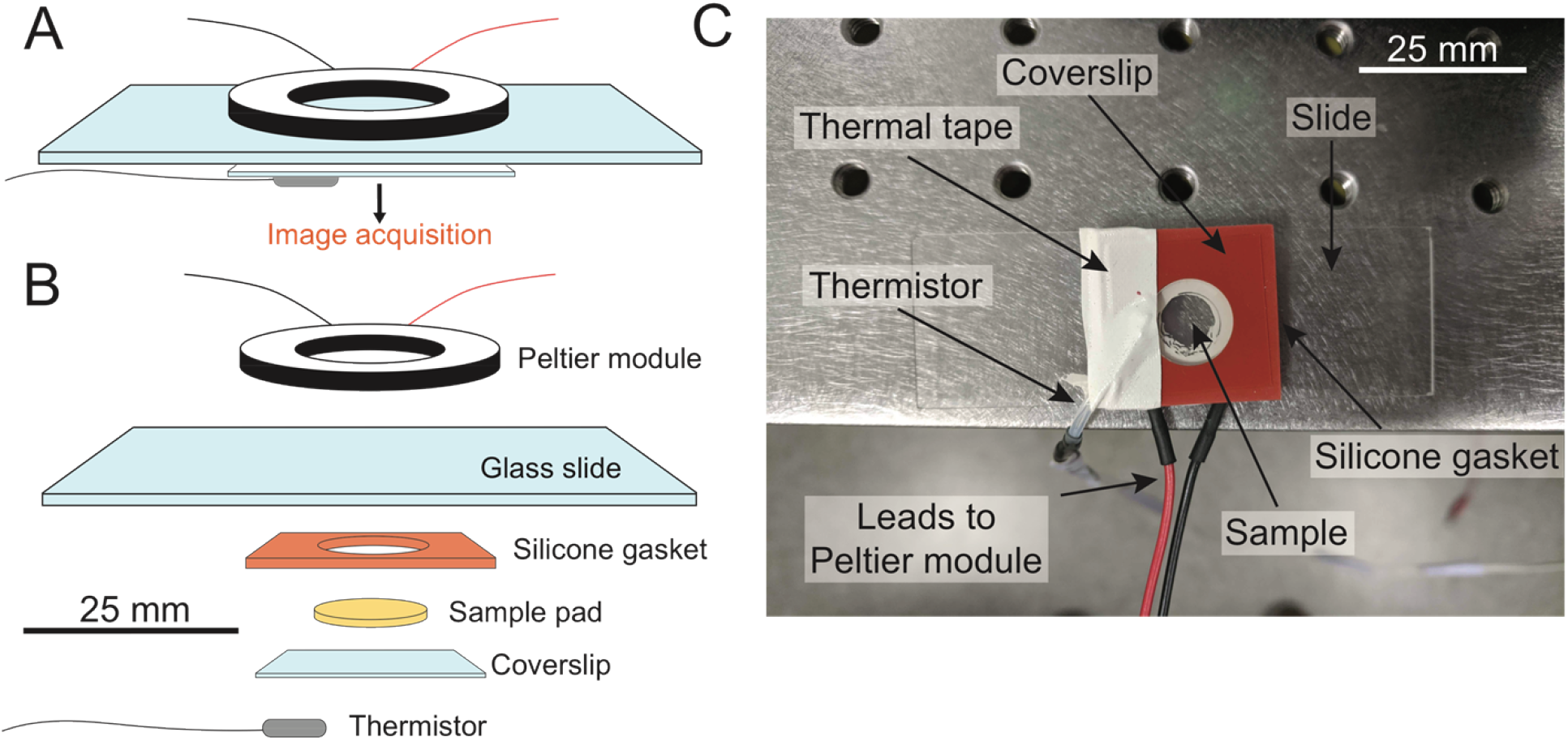
SiCTeC: an easy-to-assembly temperature controller for single-cell imaging. A) Schematic of a temperature-controlled sample slide. A ring-shaped Peltier module delivers heat to the sample, and the temperature is monitored at the coverslip using a thermistor. Images of cells can be acquired using most microscopy systems, and processed using automated segmentation and tracking algorithms. B) Exploded schematic of the slide in (A). C) Photo of sample slide with Peltier module and thermistor. The thermistor is sealed to the coverslip with thermal tape, and power is delivered to the Peltier module using electronic control components (not shown).

We monitor temperature using a thermistor, a device for which log(resistance) depends approximately linearly on the inverse of temperature, which we calibrated using a commercial thermocouple device (Methods). We sealed the thermistor to the coverslip by wrapping thermal tape around the slide and coverslip (Fig. 1C).

To actively control the temperature, we use the open-source Arduino platform to implement a proportional-integral-derivative (PID) algorithm, which is commonplace in systems design control [21] (Fig. 2A). Two separate methods control the temperature setpoint: (1) manual control, using a potentiometer with a set range of temperature values (Fig. 2B, S1A), or (2) programmed control with an executable function using the Arduino software. Having two options allows the user to either set target temperatures dynamically or to execute more complex temperature programs. The open-source software Processing [22] displays the temperature readout from the Arduino in real time and writes power and temperature data to a table (Fig. S1B). The temperature is read every 500 ms, and the error, defined as the difference between the current and target temperatures, is used as input for the PID algorithm (Fig. 2A). Starting from a 12 V power supply, we use a buck converter to step down the power input, given the 5 V limit on the Peltier module (Fig. 2B, S1A). To modulate the power, we use a motor driver, which receives a pulse-width-modulation (PWM) signal from the Arduino’s PID output (Fig. 2B, S1A). Together, these components are soldered onto a breadboard and placed into a plastic enclosure (Fig. S1A).

**Figure 2:**
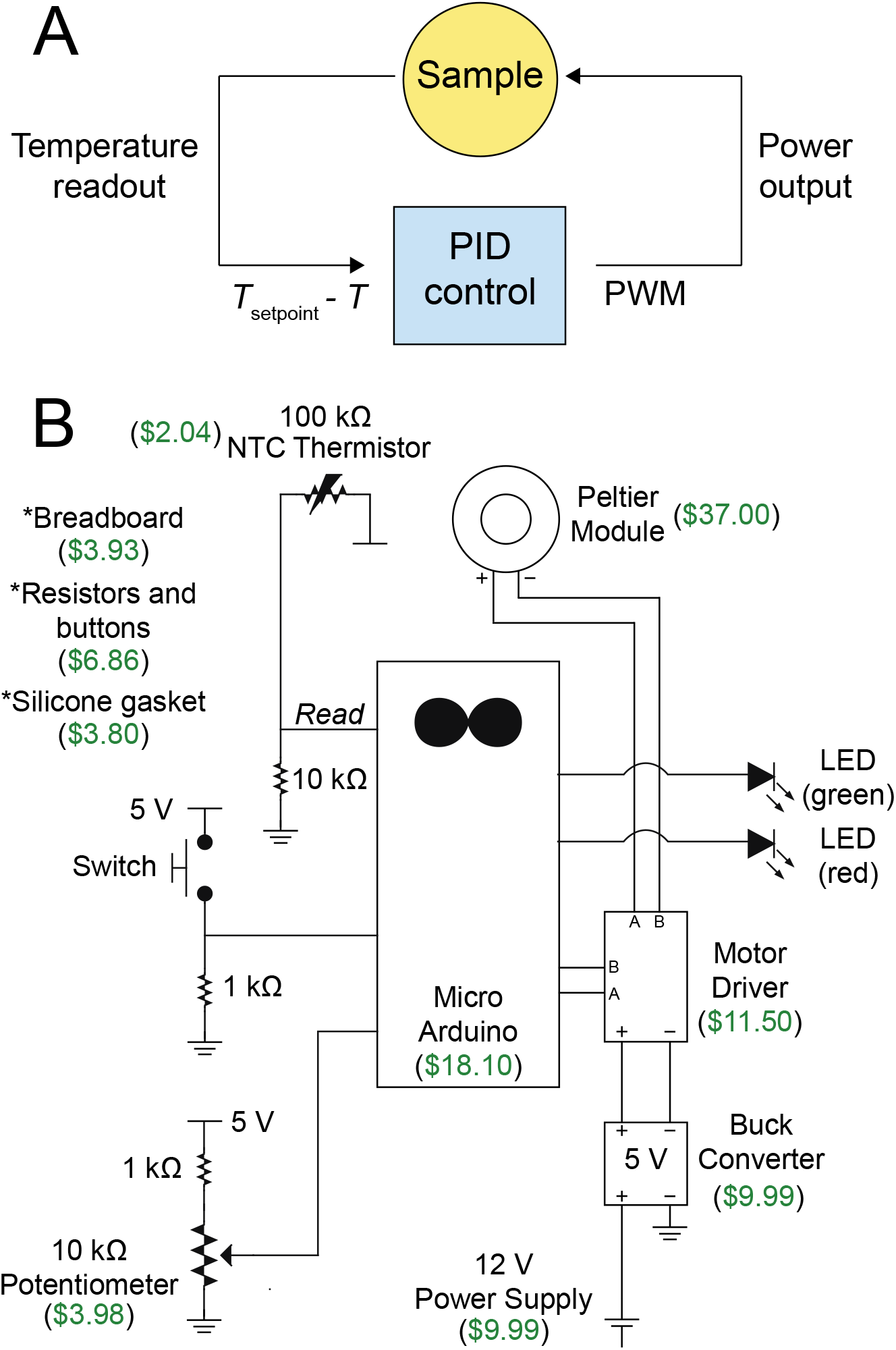
Design of an inexpensive control system. A) Control design for proportional-integral-derivative (PID) temperature control. The temperature (*T*) is read by the thermistor at the coverslip, and its difference with respect to the setpoint (*T*_setpoint_) is used as an input to the PID algorithm. Pulse-width-modulation (PWM) signals are used to modulate the power input to the Peltier module. B) Circuit design for the temperature-control system. Components are labeled with their estimated itemized price in US dollars. Asterisks indicate items not shown. The total cost is ~$107 (Table S1).

Due to their small size, imaging bacteria is conventionally performed with high-magnification oil-immersion objectives, which act as heatsinks when in contact with samples. This issue can lead to incorrect temperature measurements and/or lack of temperature control at the sample plane due to heat dissipation, and also dramatically limits the speed of temperature shifts. Further, the physical constraints of keeping the thermistor in contact with the coverslip near the sample could obstruct imaging. To circumvent these issues, SiCTeC uses a 40X air objective with a relatively high numerical aperture (NA: 0.95) and a 1.5X tube lens to achieve an effective magnification of 60X. We found that this setup produces measurements of cellular dimensions with similar precision and accuracy to a 100X oil-immersion objective (NA: 1.45) (Fig. S2).

The largest design challenge was to achieve imaging stability at high temperatures; hydrogel melting and subsequent focus and sample drifting rendered this goal problematic. To address these issues, we used a silicone gasket to secure hydrogels of at least 3% agarose (w/v), which have higher melting point and stiffness than lower agarose concentrations [23] (Fig. 1B,C). We also limited the maximum power delivered to the device at PWM = 50, out of a possible PWM = 255. In addition, we found that slower PID parameters allowed the hydrogel to equilibrate gradually, while still reaching the target temperature over a time period much shorter (1-2 min) than the doubling time of a fast-growing species such as *E. coli* (20-60 min) (Table S2).

### SiCTeC maintains and increases temperature quickly across a wide range of temperatures

To determine whether SiCTeC could maintain temperature for long periods of time, we set up an agarose pad with exponentially growing wild-type *E. coli* MG1655 cells from a 37 °C liquid culture to mimic a typical time-lapse imaging experiment and attempted to maintain the temperature of the pad at 37 °C using our device. Monitoring the temperature at the coverslip using a thermistor for 24 h revealed that the temperature was highly stable, with a mean of 37.0±0.15 °C and a maximum deviation of 0.6 °C (Fig. 3A, S3A). To validate that the temperature of the cells was essentially constant, we monitored cell growth for the first 70 min before the population became crowded, resulting in growth inhibition by surrounding cells (Fig. 3B,C). Cell morphology was maintained throughout the 70 min (Fig. 3B) and was consistent with previous studies performed at steady state [24]. Growth rate was maintained with an average of 1.98 h^−1^ (Fig. 3C), similar to previous steady-state measurements of growth at 37 °C [25], indicating that the cells experienced a constant temperature environment.

**Figure 3:**
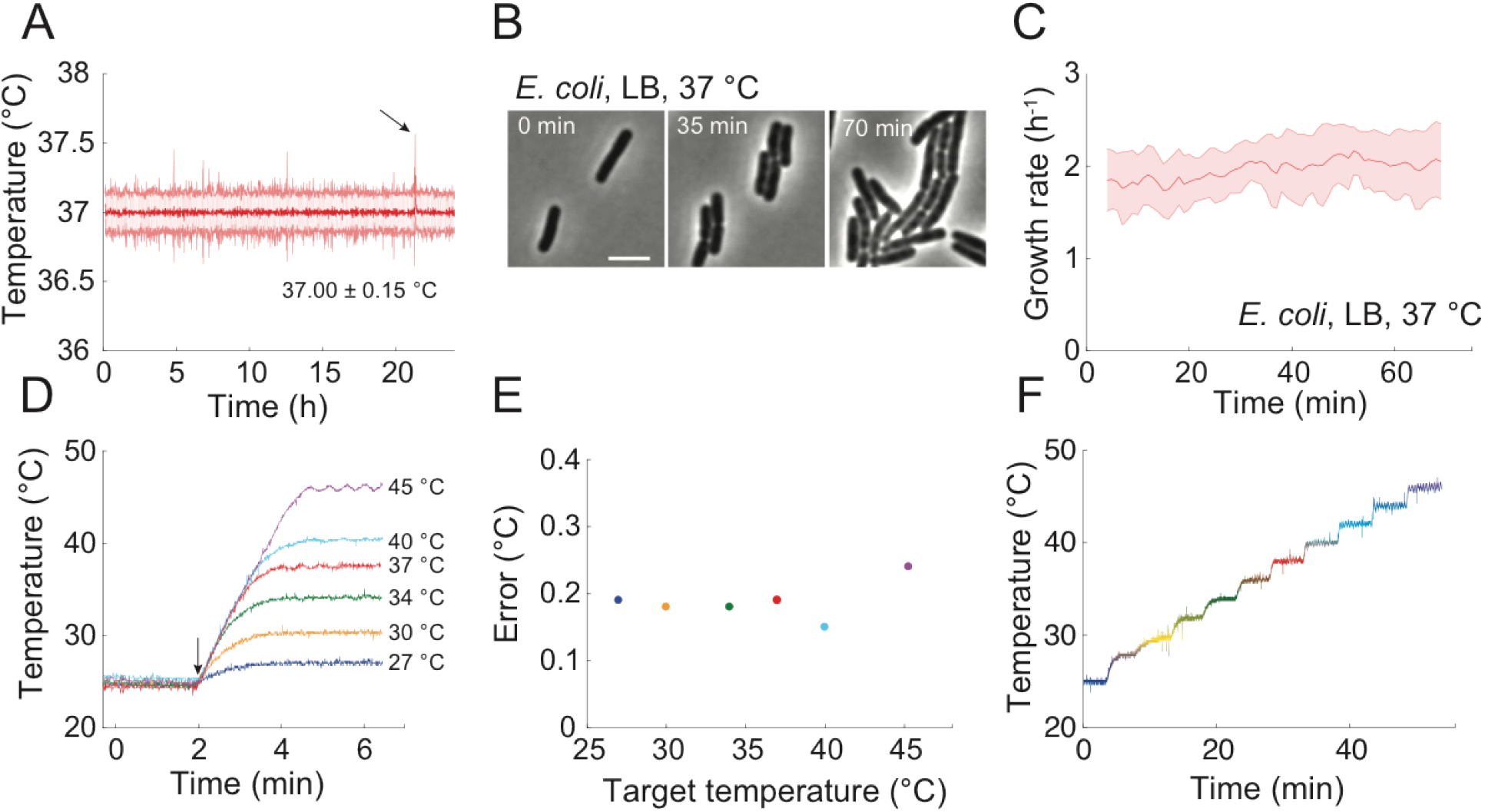
SiCTeC can maintain a wide range of temperatures with very small fluctuations. A) Temperature readout during maintenance of a sample at 37 °C for 24 h. The temperature was binned every 1 min; the shaded region represents ± 1 standard deviation. Arrow points to the most extreme outlier (0.6 °C). Time-lapse images of *E. coli* grown on an agarose pad maintained at 37 °C using SiCTeC as monitored in (A). Cell morphology and viability were maintained throughout 70 min of tracking. LB, lysogeny broth. Scale bar: 4 µm. B) Population-averaged growth rate of *E. coli* cells was approximately constant during 70 min of maintenance at 37 °C as monitored in (A). Shaded region represents ± 1 standard deviation (*n* = 301 cells). Temperature-shift experiments from 25 °C demonstrate rapid equilibration at target temperatures of 27-45 °C. The device was maintained at 25 °C for 2 minutes and then shifted to the target temperature. Arrow indicates time of shift. C) The error in temperature maintenance was less than 0.3 °C at each target temperature in (D), defined as the standard deviation over 3-min windows. D) Temperature stepping experiment from 25 °C to 46 °C demonstrates the ability of SiCTeC to achieve near arbitrary upshift waveforms. The initial step was to 28 °C, while all subsequent steps were increments of 2 °C at 5-min intervals.

To measure the capacity of our device to maintain temperature across a range of temperatures, we heated an agarose pad from room temperature to temperatures ranging from 25 °C to 45 °C (Fig. 3D). In each case, the temperature equilibrated at the desired value within 2 min, and was maintained thereafter with a standard deviation <0.2 °C (Fig. 3D,E). Up to 40 °C, the temperature control error did not depend on the target temperature (Fig. 3E); at higher temperatures the standard deviation remained low (<0.3 °C; Fig. 3E), although the slower PID parameters produced small oscillations around the target temperature (Fig. 3D), likely due to rapid cooling of the device at higher temperatures.

To further interrogate the speed and flexibility of SiCTeC, we increased the temperature from 25 °C to 46 °C in a stepwise manner (Fig. 3F). The setpoint was first increased to 28 °C and then increased by 2 °C every 5 min. In each case, the target temperature was reached within 2 min and remained stable thereafter (Fig. 3F). Higher temperatures underwent small oscillations (Fig. 3D), consistent with the single shift from room temperature to 45 °C (Fig. 3D).

SiCTeC can also achieve much higher temperatures than have been previously been considered accessible for single-cell time-lapse imaging. At ~1/5 the maximum power of the Peltier module, the device stably reached 89 °C (Fig. 4A). Shifting the temperature from 37 °C to 89 °C melted the agarose hydrogel, but the silicone gasket provided a reservoir for the liquid to be retained during imaging (Fig. 4A,B). Some of the cells remained adhered to the coverslip at high temperatures, allowing single-cell tracking throughout the 30 min at 89 °C and subsequent cooling (Fig. 4B). At 89 °C, cells maintained their shape but decreased in size, coincident with blebbing (Fig. 4B,C). Similar blebbing has been observed at very high temperatures using electron microscopy, and has been attributed to loss of outer-membrane material through vesiculation [26]. After cooling to room temperature, cells eventually became phase-bright, indicative of extreme protein stress (Fig. 4D) [27].

**Figure 4:**
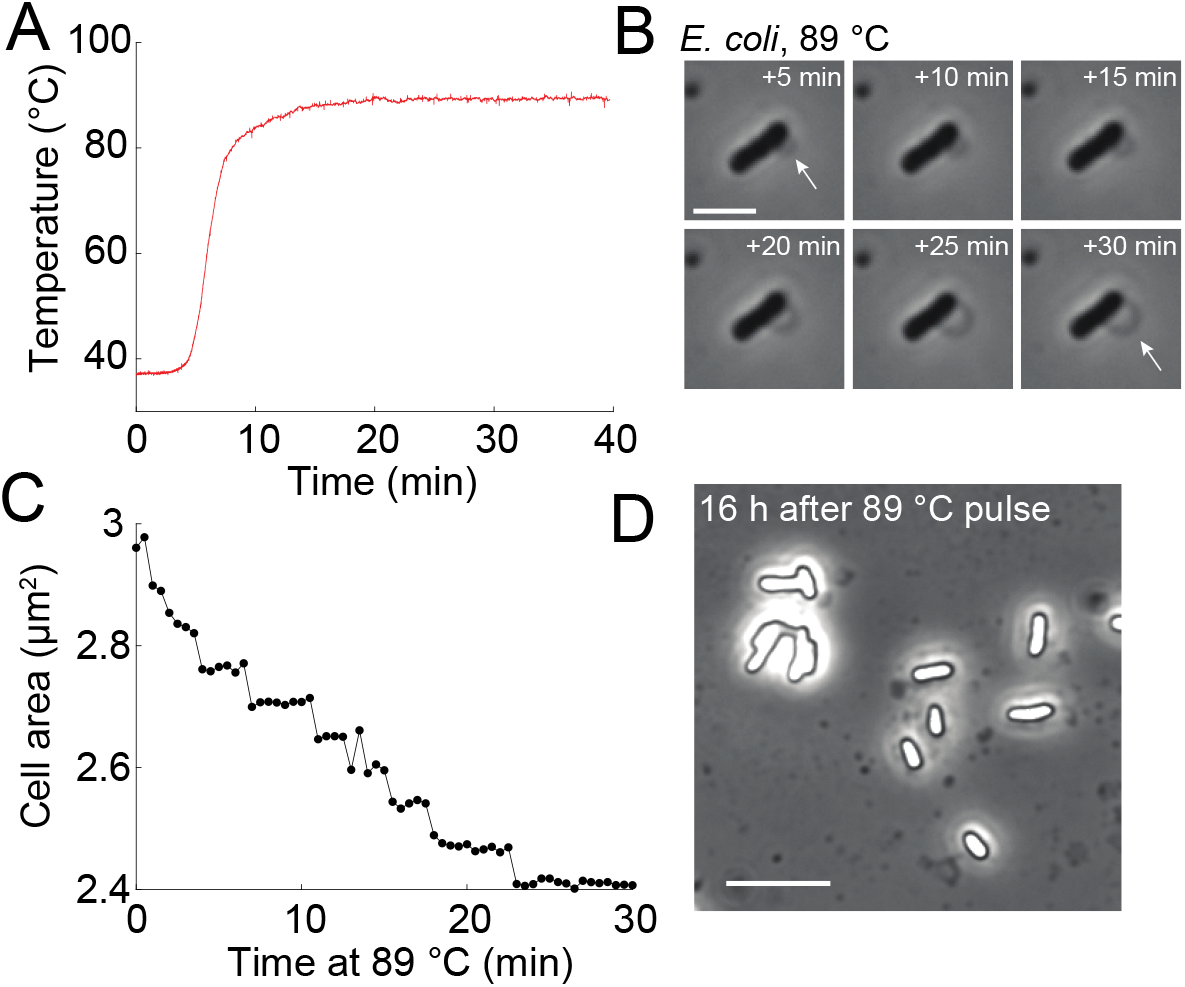
SiCTeC enables extended imaging at thermophilic temperatures. A) Temperature of an *E. coli* sample undergoing a shift from 37 °C to 89 °C, the maximum temperature allowed by the device at PWM = 50. The system achieved a steady state after 7 min. Scale bar: 3 µm. B) Time-lapse imaging of an *E. coli* cell at 89 °C for 30 min revealed halting of growth and blebbing (arrows). C) The cross-sectional area of the cell in (B) shrank progressively while temperature was maintained at 89 °C. D) Phase-contrast image of cells 16 h after a 30-min pulse at 89 °C reveals phase-bright interior indicative of the unfolded protein response [27]. Scale bar: 8 µm.

Taken together, these data indicate that SiCTeC can impose temperature increases with almost arbitrary waveforms over an extremely wide range of temperatures.

### SiCTeC enables temperature oscillations via passive cooling for rapid temperature downshifts

Complex temperature dynamics can also involve temperature downshifts. To determine the speed at which the temperature drops back to ambient levels after the device was turned off, we grew *E. coli* cells to steady state at 30 °C and then placed them on an agarose pad subjected to oscillatory cycles of upshifts to 37 °C for 10 min followed by passive cooling (PWM=0, Fig. S3B) for 10 min (Fig. 5A). Active heating to 37 °C was achieved with a time constant of ~0.6 min, similar to our previous measurements (Fig. 3D). Cooling exhibited a time constant of ~1.9 min (Fig. 5B), which is slower than the heating time constant but much faster than the doubling time of *E. coli* (~20 min at 37 °C). Thus, temperature changes in both directions can be achieved sufficiently quickly to program repeatable, precise, and nearly arbitrary temperature dynamics.

**Figure 5:**
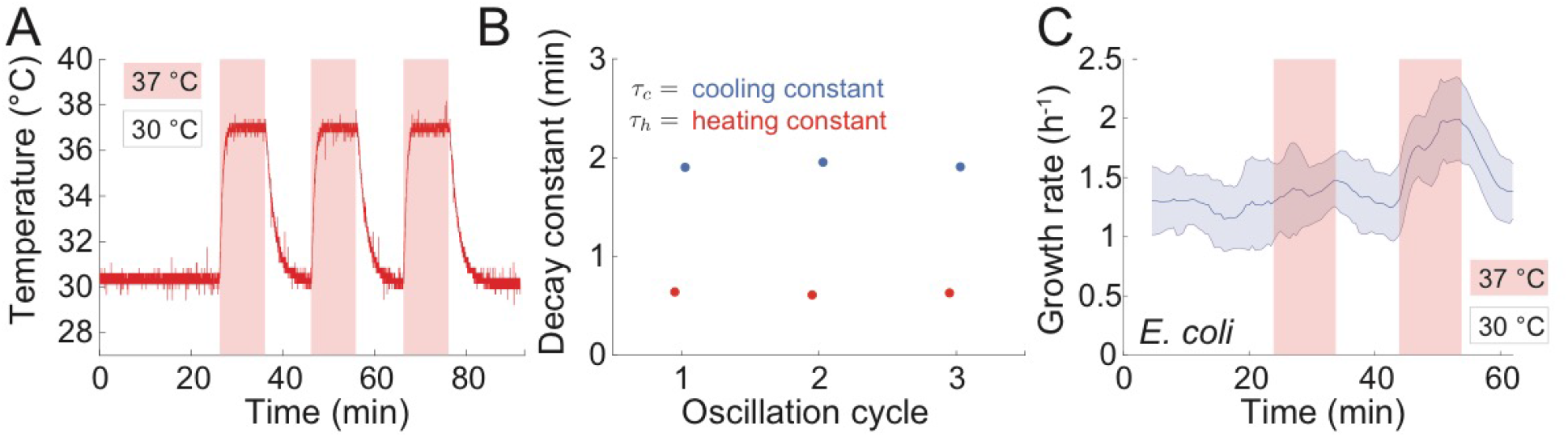
Temperature oscillations with programmable and repeatable waveforms reveal rapid adaptation of *E. coli* to temperature shifts. A) *E. coli* cells were subjected to 3 cycles of oscillations with 20-min period between 30 °C and 37 °C. Downshifts from 37 °C to 30 °C were achieved through passive cooling. The waveform remained the same shape throughout all three cycles. B) Heating and passive cooling were repeatable and rapid in each cycle in (A). The cooling constant (*τ_c_*, blue) was obtained by fitting a normalized exponential function (1-exp(-*t*/*τ*)) to each transition from 37 °C to 30 °C; the heating constant (*τ_h_*, red) was obtained by fitting the function exp(-*t*/*τ*) to each transition from 30 °C to 37 °C. Error bars (95% confidence intervals) are smaller than the data markers. C) *E. coli* cells can rapidly adapt their growth rate to temperature oscillations. Growth rate of *E. coli* cells on LB agarose pads subjected to temperature oscillations between 30 °C and 37 °C exhibited a faster increase in growth rate in the second up- and downshift cycle compared with the first cycle, indicating rapid adaptation to the oscillations. The red and blue shaded regions represent temperature upshifts and downshifts, respectively. Shaded region represents ± 1 standard deviation (*n* = 588 cells).

To determine how *E. coli* cells respond to temperature oscillations, we quantified the growth rate of hundreds of cells throughout the first 60 min of the experiment (until the pad became crowded and growth slowed). Growth rate responded slowly to the initial temperature upshift, increasing by only ~14% after 10 min (Fig. 5C). However, growth rate responded much more quickly to the subsequent upshift (Fig. 5C), reaching the steady-state value of ~2 h^−1^ within 7-8 min after the temperature increase was initiated. Downshift responses followed a similar trend: the decay of the growth rate was faster during the second downshift than during the first (Fig. 5C). These data reveal that cells adapt their response to temperature oscillations within a single oscillatory period (20 min), which is shorter than a single doubling time for *E. coli*.

### SiCTeC enables discovery of phenotypes associated with temperature-sensitive mutations

A common strategy for studying loss-of-function mutations in essential genes has been to isolate temperature-sensitive (ts) mutants that only grow at the permissive temperature [28]. In many cases, it is thought that the switch to the non-permissive temperature destabilizes the protein of interest, making it non-functional, and eventually results in growth inhibition. SiCTeC is ideal for examining how quickly growth inhibition emerges upon the temperature transition, as well as the phenotypic consequences of the switch. We focused on temperature-sensitive mutants of two penicillin-binding proteins (PBP) in *E. coli.* PBP2 is an essential enzyme responsible for cell wall cross-linking during elongation [29]. When PBP2 is fully inhibited by the antibiotic mecillinam, cells are unable to properly maintain their rod-like shape and become round [30]. PBP3 carries out cell wall cross-linking during cell division [31], and its inhibition by the antibiotic cephalexin inhibits division and causes filamentation [32]. For both proteins, temperature-sensitive mutants have been isolated in which the proteins become functionally compromised and cells ultimately cannot grow at high temperatures (42 °C) [30, 32]. SiCTeC allowed us to shift these mutants from the lower, permissive temperature (30 °C) up to the non-permissive temperature (42 °C) and to track the entire trajectory of morphological changes in individual cells; such single-cell investigations have been inaccessible using traditional techniques.

To verify single-cell phenotypes at non-permissive temperatures, we grew PBP2ts (SP4500, Methods) and PBP3ts (JE7611) cells in LB at 30 °C in liquid culture into log phase and then back-diluted the cultures and grew them at 42 °C in LB for 2 h before imaging them on agarose pads. PBP2ts cells were football-shaped (Fig. 6A, left), and PBP3ts cells were filamentous (Fig. 6A, right), as expected.

**Figure 6:**
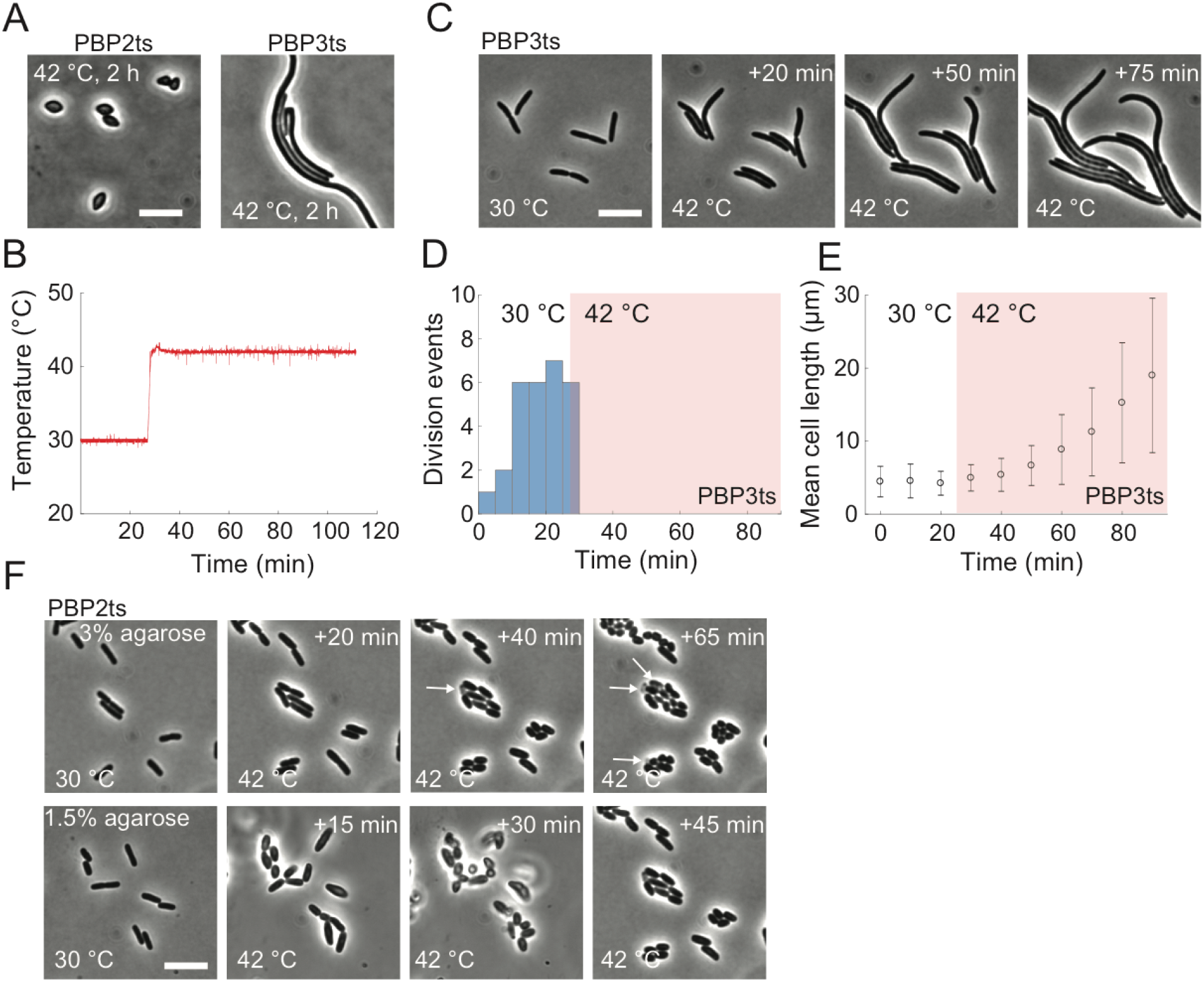
SiCTeC enables quantification of temperature-sensitive phenotypes during transitions to non-permissive temperatures. A) After 2 h of growth in liquid at the non-permissive temperature 42 °C, temperature sensitive (ts) mutants of the cell wall synthesis enzymes PBP2 (left, strain SP4500) and PBP3 (right, strain JE7611) adopted football-shaped and filamentous morphologies due to inhibition of the elongasome and divisome, respectively. Scale bar: 8 µm. B) SiCTeC supported the desired temperature profile, as indicated by the temperature readout while shifting PBP2ts cells from its permissive temperature (30 °C) to 42 °C. C) After a shift from 30 °C to 42 °C, PBP3ts cells rapidly halted division and elongated filamentously, as expected due to inhibition of cell division. Scale bar: 8 µm. D) Division events of PBP3ts cells, binned into 5-min windows, were only observed during growth at 30 °C or almost immediately after the shift to 42 °C (*n =* 33 total division events). E) Mean cell length of PBP3ts cells increased continuously throughout the shift from 30 °C to 42 °C (*n* = 31-40 cells per data point). F) PBP2ts mutants are sensitive to hydrogel stiffness. After a shift from 30 °C to 42 °C, the PBP2ts mutant grown on 3% agarose pads (top) remained approximately rod-shaped rather than adopting the rounded phenotype in (A), and a subpopulation of cells lysed (arrows). By contrast, cells grown on 1.5% agarose pads (bottom) became football-shaped as in (A) within 30 min, without noticeable cell lysis. Scale bar: 8 µm.

To monitor single-cell morphology during the upshift, we grew the PBPts mutants into steady state (Methods) in liquid LB at 30 °C, placed them on a 3% agarose pad, and shifted the temperature to 42 °C using SiCTeC (Fig. 6B). In PBP3ts cells, division halted almost immediately but cells continued to elongate (Fig. 6C,D). Within 75 min after shifting to 42 °C, mean cell length increased nearly 4-fold (Fig. 6E), while the growth rate was maintained (Fig. S4B). Thus, the temperature shift inactivates PBP3 almost immediately without causing any obvious side effects at the cellular level.

On 3% agarose pads, PBP2ts cells remained approximately rod-shaped 40 min after the shift to 42 °C (Fig. 6F, top). By 40 min, some cells had started to lyse (Fig. 6F, top); lysis continued at 100 min, at which point cells were either small spheres or short rods (Fig. S4A). These data suggested that growth on the 3% agarose pad unexpectedly altered the morphological trajectory and compromised the mechanical integrity of the cell wall. To test this hypothesis, we performed a similar experiment on 1.5% agarose pads, which have substantially lower mechanical stiffness [23]. In this case, we observed swelling of PBP2ts cells within 15 min after shifting to 42 °C, and the football-shaped phenotype was reached within 45 min without any signs of lysis (Fig. 6F, bottom).

These results demonstrate that SiCTeC enables novel biological discoveries by providing information about single-cell dynamics that is inaccessible in bulk liquid-culture experiments.

## DISCUSSION

Here, we have demonstrated that a stage-top temperature controller for single-cell imaging at above-ambient temperatures can be assembled using low-cost and accessible components (Table S1). SiCTeC can achieve relatively fast temperature increases (1-2 min; Fig. 3D), and passive cooling is sufficiently fast for single-cell studies (4-5 min; Fig. 5A,B). While the SiCTeC can in principle achieve much faster heating using more aggressive PID parameters, we found that melting of the pad and subsequent image plane and cell drifting made such conditions incompatible with traditional agarose pad-based single-cell imaging. To achieve faster cooling, the device would need to draw heat to maintain a temperature gradient (heatsink) and include a method for quick heat dissipation (such as a fan). While we were able to incorporate both features in prototype devices, our studies indicate that the simpler and less complex device we have presented here is sufficient for studies of cell growth. A large heatsink with fan-assisted heat dissipation would potentially allow for stable below-ambient temperatures in future iterations of the device.

Some commercial devices are capable of very short heating and cooling times (~10 s) using a fluidic approach [33]. These devices are limited in agarose pad sample sizes, making long-term experiments difficult. Using large agarose pads, we can image for many hours on our device without drying or starvation. Further, fluidic designs have a maximum temperature of 45 °C [33], whereas SiCTeC can maintain extreme temperatures of nearly 90 °C (Fig. 4). The flexibility of our device in terms of programming near-arbitrary temperature dynamics is also unmatched. While other modified devices have recently been constructed to allow for imaging at extreme temperatures [34, 35], these instruments are more expensive than SiCTeC, highly customized, and limited to fluorescence imaging. Our device is entirely compatible with both brightfield and fluorescence imaging, which can be utilized to study protein localization and expression changes during temperature shifts as long as the temperature dependence of fluorescence is accounted for [36].

Perhaps most importantly, SiCTeC offers a highly affordable, flexible, reusable alternative that can be constructed by nearly any lab. Our device has many use cases, such as on-the-fly temperature changes and more complex, programmable temperature changes. Sample temperature can be varied in a stepwise fashion (Fig. 3) or driven in oscillations (Fig. 5); while the consequences of both of these perturbations have yet to be fully explored, here we discovered that during temperature oscillations *E. coli* cells experience a much faster acceleration of growth during the second temperature upshift than during the first (Fig. 5C). By employing open-source components and software, this design also allows for customizable and user-specific needs. Fundamentally, this design makes a minimal alteration to the traditional (and inexpensive) agarose pad technique familiar to microbiologists, with components collectively priced at less than $110 (Table S1) and assembled in just a few hours.

For bacteria, temperature changes are unavoidable within a host (e.g. fever) and in the environment (diurnal cycles and climate change on short and long timescales, respectively). Yet, we know little about how key processes such as bacterial responses to antibiotics depend on temperature. Tracking single-cell dynamics is often critical to understanding how cells respond to environmental perturbations. For instance, *E. coli* cells grow more slowly at steady-state at high osmolarity, yet maintain their growth rate immediately after an increase in medium osmolarity [37], demonstrating that turgor pressure does not determine growth rate (despite the decrease in steady-state growth rate as osmolarity increases). In this capacity, SiCTeC has the potential to spur future studies of the temperature dependence of cellular physiology in a wide variety of organisms. Agarose pads can be used to modify the nutrient and chemical environment, and the modification of pad stiffness provides the ability to examine the impact of mechanics on cellular responses to temperature changes. Temperature can even be used to modulate the stiffness of a hydrogel [38], enabling the study of how cells adapt to dynamic substrate mechanics. With a simple extension of our design, we anticipate that a similar Peltier-based strategy can be incorporated into a microfluidic platform [39], enabling both long-term imaging (e.g. repeated oscillations) and chemical perturbations throughout a single temperature-shift experiment. The straightforward and inexpensive construction of our device should facilitate addressing many important questions across organisms that inhabit environments whose temperatures are well-regulated (e.g. the mammalian gut) and fluctuating (e.g. soil).

## METHODS

### Strains and cell culture

The *E. coli* strains used in this study were MG1655, *ftsI730* (JE7611, PBP3ts) [40], and *pbpA45* (SP4500, PBP2ts) [41]. All experiments were carried out in rich medium (lysogeny broth, LB).

Wild-type cells were inoculated directly into LB from frozen stocks and grown overnight at 37 °C, then diluted 1:200 into fresh LB and grown for 2 h at the desired temperature. For temperature-shift experiments, cells were diluted 1:10 twice at the initial temperature to establish steady-state growth.

PBPts strains were grown from frozen stocks overnight at 30 °C, diluted 1:100 into fresh LB, and grown for 4 h (due to their slower growth rates). Cells were then diluted 1:10 twice to establish steady-state growth before shifting to the non-permissive temperature.

### Thermistor calibration

From 20 °C to 50 °C, thermistor temperature *T* is approximately related to the resistance *R* through the equation 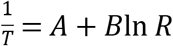. Thus, we estimated *B* based on pairwise measurements at two temperatures *T*_1_ and *T*_2_ using the equation 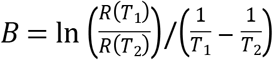. The resistances at 4, 22, 30, and 37 °C were 274, 114, 81, and 60 kΩ, respectively. The average estimate of *B* across all pairwise combinations was 3950 °C^−1^, suggesting that the nominal resistance at 25 °C was close to 100 kΩ, in agreement with the manufacturer’s specifications (https://www.makeralot.com/download/Reprap-Hotend-Thermistor-NTC-3950-100K.pdf). Measurements were in close agreement with a highly sensitive thermocouple (Omega Engineering).

### Sample preparation and imaging

LB-agarose pads were prepared by boiling 1.5% or 3% ultrapure agarose (Sigma Aldrich) in LB, then pipetting 150 μL of liquid agarose medium into a silicone gasket (Grace Bio-Labs) on a glass slide, which was compressed with another glass slide. The pad was allowed to solidify at the initial temperature for shift experiments. Typically, the microscope was kept at the initial temperature using a temperature-controlled microscope enclosure (Haison Technology).

One microliter of cells at the initial temperature was placed onto the center of the pad and allowed to dry at the initial temperature before placing a coverslip on the pad. The thermistor was sealed on the coverslip with thermal tape. The device was placed onto the microscope slide mount, and leads from the electrical control components were attached. During time-lapse imaging, the temperature readout and PWM signal were visualized in real time using the open-source software Processing v. 3.5.4. All relevant data from the device were saved as a .csv file upon exit from the visualization tool.

Samples were imaged in phase-contrast using a Ti-Eclipse stand (Nikon Instruments) with a 40X Ph2 air objective (NA: 0.95) (Nikon Instruments) along with a 1.5X tube lens, or a 100X Ph3 oil-immersion objective (NA: 1.45) (Nikon Instruments). Images were acquired using a Zyla 4.2 sCMOS camera (Andor Technology); time-lapse images were acquired every 30 s. The microscope system was integrated using μManager v. 1.41 [42].

### Image analysis

To account for the drift that can occur during temperature shifts (data not shown), time-lapse data were aligned using the Template-Matching plugin [43] in ImageJ [44]. These aligned images were segmented by the deep neural network-based machine learning framework *DeepCell* [45]. To train the network, approximately 200 cells were manually outlined to produce a training dataset for our specific imaging conditions. Trained networks were used to generate binary images for feature (extracellular/cell perimeter/cytoplasm) identification. These images were used as the input for gradient segmentation in *Morphometrics* v. 1.1 [46] to define cell contours at sub-pixel resolution. Custom MATLAB (Mathworks) scripts were used to track individual cells, and to measure the centerline of rod-shaped cells to calculate cellular geometries [47]. Growth rate was defined as the time-derivative of the logarithm of cell length, and growth-rate trajectories were binned to show population-average behaviors.

## Author Contributions

B.D.K., L.Z., and K.C.H. designed the research; B.D.K. and L.Z. performed the research;

B.D.K. and L.Z. analyzed the data; and B.D.K., L.Z., and K.C.H. wrote the manuscript.

## Acknowledgments

The authors thank members of the Huang lab and George Korir for helpful discussions, and Paul Rujigrok, Heidi Arjes, and Fred Chang for comments on the manuscript. The authors acknowledge funding from the Allen Discovery Center at Stanford on Systems Modeling of Infection (to K.C.H). K.C.H. is a Chan Zuckerberg Biohub Investigator.

